# Wind speed affects the rate and kinetics of stomatal conductance

**DOI:** 10.1101/2023.12.06.569782

**Authors:** Or Shapira, Uri Hochberg, Scott McAdam, Tamar Azoulay-Shemer, Craig R. Brodersen, Noel Michelle Holbrook, Yotam Zait

**Affiliations:** Golan Research Institute, P.O. Box 97, Katzrin 12900, Israel; Fruit Tree Sciences, Agricultural Research Organization (ARO), The Volcani Center, Newe Ya’ar Research Center, Ramat Yishay, 30095, Israel; ARO Volcani Center, Institute of Soil, Water and Environmental Sciences, Rishon Lezion, 7505101 Israel; Purdue Center for Plant Biology, Department of Botany and Plant Pathology, Purdue University, West Lafayette, IN 47907, USA; School of the Environment, Yale University, New Haven, CT 06511, USA; Department of Organismic and Evolutionary Biology, Harvard University, 26 Oxford St, Cambridge, MA, 02138 USA; The Robert H. Smith Institute of Plant Sciences and Genetics in Agriculture, Faculty of Agriculture, Food and Environment, The Hebrew University of Jerusalem, Rehovot 76100, Israel

**Keywords:** Transpiration, Stomata, gas exchange, wind, boundary layer, leaf fan

## Abstract

Understanding the relationship between wind speed and gas exchange in plants is a longstanding challenge. Our aim was to investigate the impact of wind speed on maximum rates of gas exchange and the kinetics of stomatal responses. We conducted experiments using an infrared gas analyzer equipped with a controlled leaf fan, enabling precise control of the boundary layer conductance. We first showed that the chamber was adequately mixed even at extremely low fan speeds (down to 200 rpm, equivalent to a wind speed of 0.0005 m s^-1^) and evaluated the link between fan speed, wind speed, and boundary layer conductance. We observed that higher wind speeds led to increased gas exchange of both water vapor and CO_2_ in Arabidopsis, presumably due to its effect on transpiration and the consequential reduction in epidermal pressure that led to stomatal opening. We documented that stomatal opening in response to light was three times faster at a fan speed of 10000 rpm (wind speed of 2 m s^-1^) compared with 500 rpm (0.25 m s^-1^) in *Vicia faba*, while the latter exhibited an opening rate that was similar to those of epidermal peels. The increase of stomatal conductance under high wind was observed in four species under field conditions. Our findings demonstrate the importance of the size of the boundary layer on determining maximum rates of gas exchange and the kinetics of gas exchange responses to environmental changes.

## Introduction

Stomatal pores play a critical role in regulating gas exchange between a plant and its environment, affecting both photosynthesis and transpiration. The aperture of stomatal pores is influenced by various environmental factors, including light, humidity, temperature and atmospheric CO_2_ (Assmann & Jegla, 2016; Kim et al., 2010; Kollist et al., 2014; Shimazaki et al., 2007). It has been previously suggested that wind influences transpiration through a direct effect on the boundary layer (Aphalo & Jarvis, 1993; Foster & Smith, 1986), but the wind effect that is mediated through stomatal regulation is far less explored and is absent from transpiration models.

Wind is hypothesized to affect gas exchange in two ways. First, wind speed affects the boundary layer, a thin layer of air adjacent to the leaf surface, where the gas flow is dominated by shearing forces, resulting from the interaction between the leaf and the surrounding air (Cowan, 1978). The thickness of this boundary layer is mainly influenced by local wind speed and leaf size, with leaf shape having a secondary effect (Nobel, 2020). The thickness of the boundary layer impacts transpiration, as it determines the resistance to water vapor diffusion from the stomatal pores to the surrounding atmosphere (Nobel, 2020). Low wind speeds lead to a thick boundary layer, and its resistance could become dominant with respect to transpiration in winds lower than 0.25 m s^-1^ (Foster & Smith, 1986). Absent or very low speed winds are relatively common in dense canopies of forests (Renaud et al., 2011) or crops (Shaw, 1977). Plants can indirectly control boundary layer conductance through modifications to morphology, size, and leaf orientation, which in turn affects flow patterns. While the variability within canopies and among species could be substantial, the interaction between wind speed, the boundary layer, and its direct effect on transpiration is well accepted (Nobel, 2020).

The second effect, which is the focus of this current research, is far less explored. Swift changes to boundary layer conductance (g_b_) caused by altered wind speed has the potential to result in rapid changes to leaf evaporation rate and thus leaf water status. Stomatal responses to changes in transpiration rate can be both actively and passively regulated (Franks, 2013). Active stomatal responses are a function of ion pumping or efflux, resulting in changes in guard cell osmotic potential (Kearns & Assmann, 1993). Passive movement, on the other hand, is a faster response that occurs as a consequence of changes in leaf water status (Buckley, 2005; McAdam & Brodribb, 2014; Meidner & Heath, 1963; Zait et al., 2017). In angiosperms the passive response is governed by the turgor of the epidermal cells which have a mechanical advantage over the guard cells (DeMichele & Sharpe, 1973; Mott & Franks, 2001; Buckley *et al*., 2011). If a rapid increase in transpiration rate causes turgor pressure to decrease in both the epidermal cells and guard cells (Franks et al., 1998), the stomata will open (Franks & Farquhar, 2007), resulting in a passive stomatal opening (Frensch & Schulze, 1988; Raschke, 1970). This passive mechanism also closes the pore when transpiration rate decreases rapidly and epidermal backpressure increases (Zait et al., 2017). We do not know the effect g_b_ on potential changes to epidermal cell turgor and thus stomatal sensitivity to environmental changes. There is very little work that has investigated whether epidermal cell turgor alters stomatal sensitivity to environmental changes (Franks and Farquhar 2007).

In this study, we investigate the relationship between wind speed and transpiration to disentangle the effects of the boundary layer on gas exchange. We relate wind speed inside the gas exchange chamber to g_b_ and examined stomatal responses to light under different wind speeds. We hypothesize that in angiosperms, increasing transpiration by increasing wind speed and thus decreasing g_b_ will result in both a passive increase in stomatal conductance and an increase in the rate of stomatal opening in response to light.

## Materials and Methods

We performed three experiments using the LI-6800 (LI-COR Biosciences, Lincoln, NE, USA). The first experiment was designed to test whether under a very low fan speed there is sufficient mixing to accurately measure gas exchange. In the second experiment, we linked the fan speed to wind speed using an omnidirectional air velocity transducer and determined g_b_ by determining evaporation from wet filter paper inside the chamber. The third set of experiments was conducted on *Vicia faba* and Arabidopsis, under controlled conditions, and four angiosperm species growing outside, to evaluate the effect of different wind speeds on plant gas exchange.

### Plant material and growth conditions

The *Vicia faba* and Arabidopsis plants used in this study were grown under controlled environmental conditions. *Vicia faba* were planted in 5 L pots, growth chamber maintained at a temperature range of 22-25°C during the day and 18-20°C at night, under a daytime light intensity of 300 μmol m^−2^ s^-1^. The relative humidity within the growth chamber was maintained at 60-70% to ensure adequate moisture availability for the plants and prevent excessive transpiration. Arabidopsis (Columbia, Col-0) seeds were planted in 250 ml pots filled with a soil mixture (Green 20, Even Ari, Israel) + 2 g/L Osmocote. Plants were grown in a growth chamber under a light intensity of 250 μmol s^−1^ m^−2^ and a 12 h photoperiod. The temperature was maintained at 21°C during the day and 19°C at night, with a relative humidity (RH) ranging between 60% during the day and 85% at night.

To test the response to wind speed in plants that are growing outdoors, and have thus experienced a far more diverse wind regimes, we measured four plant species growing at Zemach Nisyonot research farm on experimental plots: mango (*Mangifera indica*), papaya (*Carica papaya*), *Withania somnifera*, and fig (*Ficus carica)*. All the measured plants were fully watered and in a healthy state.

### Gas exchange measurements

Gas exchange measurements were conducted using a LI-6800F portable photosynthesis system (LI-COR Biosciences, Lincoln, NE, USA). This system is equipped with a leaf chamber of 2 cm^2^ and an infrared gas analyzer (IRGA) to measure CO_2_ assimilation rate, transpiration rate (E), and other climatic parameters in real-time. On the day of the experiment, a healthy, fully expanded leaf was selected for measurements. The leaf was carefully inserted into the leaf chamber, and the system was set to control all environmental parameters including light intensity, temperature, relative humidity, and CO_2_ concentration (see details below). The leaf fan speed was adjusted in the chamber to manipulate boundary layer conductance during the experiment.

### Evaluation of mixing in the LI-6800F chamber

In this experiment, we employed a methodology that closely followed the approach of McNab (2006) for measurements of animal respiration. A mature mango leaf, attached to the plant, was placed into a LI-COR 6800F cuvette and subjected to an hour of dark adaptation until a stable dark respiration rate was achieved. We then set the flow rate to 950 µmol s^-1^ for five minutes, recording the respiration rate every five seconds. Following this, we reduced the flow rate to around 850 µmol s^-1^ for an additional five minutes. This procedure was repeated at ten lower flow rates, down to 20 µmol s^-1^. The procedure was repeated at four fan speeds (200, 800, 2000, and 10,000 rpm), with the aim of observing a reciprocal linear relationship between respiration rate and flow rate, to verify adequate air mixing within the chamber (McNab, 2006). Any deviation from a linear relationship between respiration rate and either fan speed or flow rate would suggest inadequate gas mixing. We identified the critical flow rate for each fan speed, defined as the minimum flow rate necessary for comprehensive gas mixing in the chamber.

### Wind speed measurements

Wind speed was measured using an omnidirectional air velocity transducer (model 8475, TSI, Singapore) placed inside the leaf chamber and connected directly to an auxiliary channel trough the 25-pin connector on the of the LI-6800F. The voltage from the sensor was transformed to wind speed in m s^-1^ according to the user manual. The data was logged into the gas exchange results file. Fan speed was changed from 200 ppm up to 10000 rpm at 200 rpm increments for 4 min at each speed. The data was logged every 15 s. The mean of the last 15 observations from each step were averaged. This test was repeated nine times at different angles of the sensor inside the measuring chamber, and averaged. The measuring probe was directed to be in the center of the 2 cm^2^ round chamber parallel to the leaf plane. Flow rate was adjusted to 500 µmol s^-1^. The area around the sensor rod and the leaf chamber interface was sealed with dental epoxy to prevent leaks.

### Measurement and calculation of boundary layer conductance

Rates of water loss from filter paper (Whatman no.3) saturated with distilled water have been used to calculate the boundary layer conductance for water vapor (*g*_*b*_) (Parkinson, 1985). This experimental approach provides a means to estimate g_b_ of unadorned leaves under the conditions inside the chamber without the interference of stomata. Chamber temperature (T_exchange_) was set to 22 °C, T filter paper was 19.7±0.9 °C, RH= 50%, VPD∼0.74±0.09 kPa, and the flow rate was 630 µmol s^-1^. The leaf thermocouple was touching the filter paper. Fan speed was changed from 0 to 300 rpm and then to 10000 rpm in increments of 200 rpm, with 4 min at each fan speed. Evaporation rate was logged every 4 s. The H_2_O IRGA was matched every 5 min and points around the match event were excluded from the results. The boundary layer ;proportion of total conductance (g_t_) was calculated by the LI-6800F according to the equation: 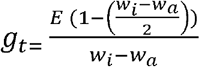 where E is the transpiration and *w*_*i*_ is the water saturation in the wet filter paper and the *w*_*a*_ is the water concentration in the air.

### Evaluation of boundary layer conductance and wind speed effects on gas exchange measurements

To examine the effect of fan speed on transpiration (E) *g*_*s*_, steady-state measurements of *g*_*s*_ and E were conducted on 6-week-old Arabidopsis plants. The leaves were stabilized in the chamber under 60% RH, T_exchange_ = 22°C, PAR 350 µmol m^-2^ s^-1^, flow rate 530 µmol s^-1^ and fan speed of 1000 rpm (wind speed of 0.03 m s^-1^). Data was recorded every 30 s. The fan speed was then increased to 10000 rpm (wind speed of 2 m s^-1^). These measurements were performed at a low CO_2_ concentration of 100 ppm to ensure stomata were open and reduce any effect of increasing internal CO_2_ concentration on stomatal movements.

Gas exchange measurements were also carried out to assess the impact of gradual changes in wind speed. The wind speed was gradually (linearly) increased over a 5-min interval by incrementally adjusting the fan speed from 200 rpm (corresponding to 0.005 m s^-1^) to 7000 rpm (equivalent to 1.5 m s^-1^). Values were logged every 30 s. Other conditions in the chamber were as described in the previous paragraph.

To assess the influence of wind speed on the kinetics of stomatal opening in the transition from dark to light, we conducted experiments with three different fan speeds (500, 1000, and 10000 rpm) while maintaining the plants in dark conditions and subsequently exposing them to light at an intensity of 800 µmol m^-2^ s^-1^. Stomatal conductance was measured throughout this transition, and we compared the rates of change in gas exchange (stomatal conductance) with the stomatal aperture observed in epidermal peels submerged in a buffer solution derived from the same leaves that were measured (see below the stomatal aperture assay). To facilitate comparison, we converted the data to a percentage of stomatal opening [(gs max-gs)/ gs max].

In the common garden experiment, measurements were conducted under the following chamber environmental conditions: a flow rate of 700 µmol s^-1^, PAR 2000 µmol m□^2^ s□^1^, a temperature of 36°C, and a relative humidity of 45%. We tested every species when wind speed was increased in one step change from 0.05 m s^-1^ to 1.5 m s^-1^ and to 2.5 m s^-1^

### Stomatal aperture assay

Fully expanded *Vicia faba* leaves were harvested from 4-week-old plants grown under controlled environmental conditions. These plants were the same ones used for gas exchange measurements with the LI-6800F system. Epidermal peels were prepared using a gentle peeling technique to ensure the integrity of the stomatal complexes (Zhu et al., 2016). Initially, the epidermal peels were incubated in darkness for 1.5-2 hours in a buffer solution containing 5 mM KCl, 1 mM CaCl_2_, and 10 mM MES-KOH (pH 6.15). After this incubation period, the peels were transferred to the light (400 µmol m^-2^ s^-1^) with 50 mM KCl and 0.1 mM CaCl_2_. Stomatal apertures were then measured using a light microscope ECHO (Rebel, Bico Company, San Diego, USA)

## Results

### Evaluation of mixing in the LI-6800F chamber

We first sought to determine the minimum fan speed in the cuvette of the gas analyzer that could provide sufficient mixing so that gas exchange could be accurately measured. We assessed the mixing inside the LI-6800F chamber by measuring leaf respiration in the dark across flow rates and fan speeds (**Fig. 1**). Because dark respiration is independent of chamber flow rate and fan speeds under adequate air mixing conditions within the chamber, a reciprocal linear relationship should exist between flow rate and ΔCO_2_. This quantitative reciprocity defines the range of flow rates (at different fan speeds) for which gases in the chamber are sufficiently mixed, ensuring that the calculated rates of gas exchange are reliable estimates. To investigate the impact of fan speeds on air mixing within a leaf chamber, the flow rate was adjusted from ∼950 to ∼50 µmol s^-1^, while maintaining constant fan speeds at 200, 800, 2000, and 10000 rpm (**Fig. 1**). We monitored the differences in CO_2_ mole fraction (µmol) under these conditions. The results revealed a significant effect of the flow rate on the curvature patterns and standard deviation associated with CO_2_ mole fraction differences. Within the observed data, a distinct linear region represented the range where reliable measurements of gas exchange could be obtained, indicating adequate air mixing. However, critical flow rates were identified, beyond which air mixing was compromised, leading to nonlinearity in the relationship. The slope of the relationship between flow rate and CO_2_ differential remained consistent across tested fan speeds, (**Figure S1)**.

**Figure 1:**
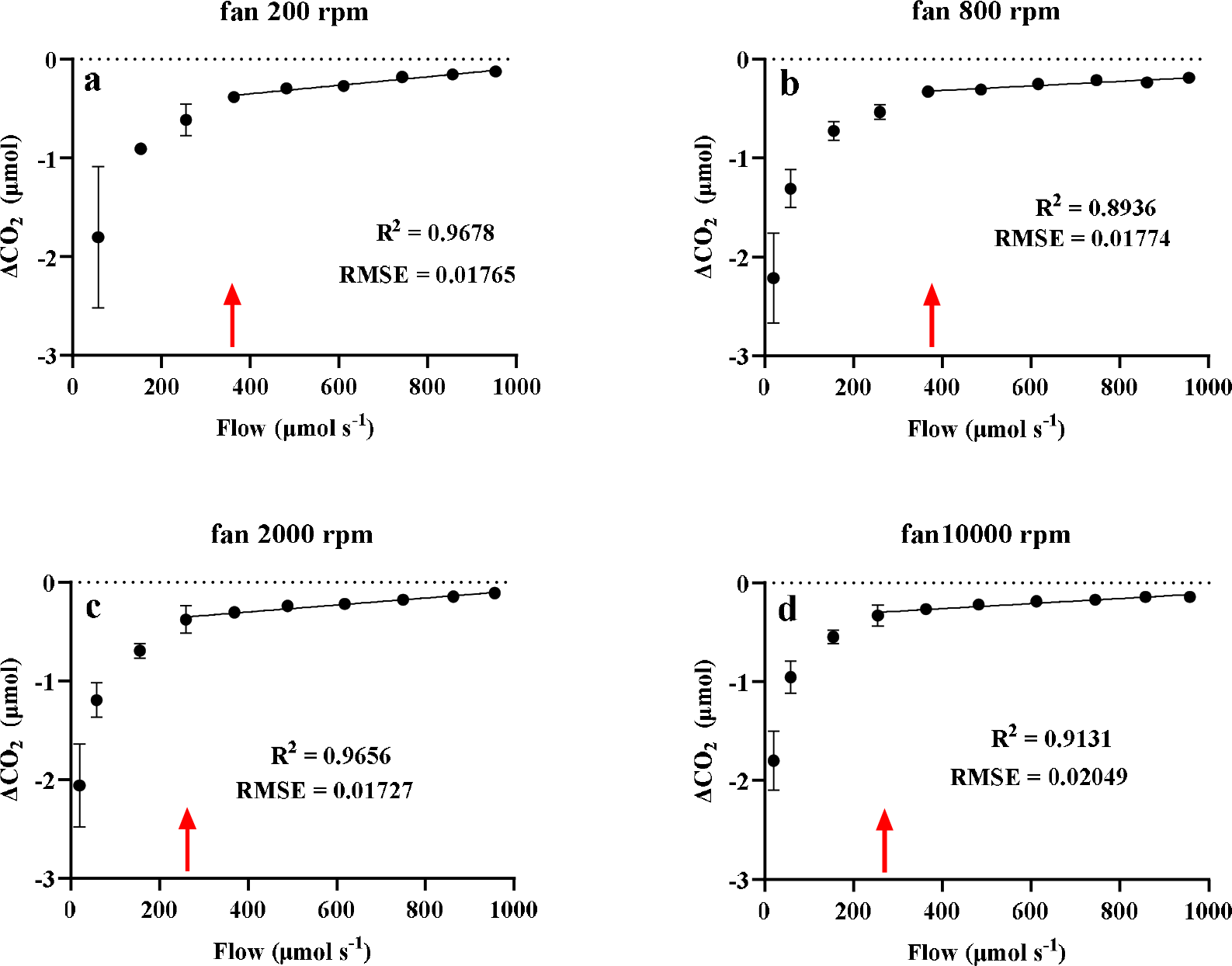
Relationship between flow rate and CO_2_ differential (respiration in the dark from a mango leaf) inside the LI-6800 leaf chamber at four different fan speeds: 200 rpm (a), 800 rpm (b), 2000 rpm (c), and 10000 rpm (d). The line represents the linear relationship between delta CO_2_ and flow rate where the chamber air mixing is above the critical flow rate point. The red arrow represents the anticipated critical point below which air mixing is not sufficient. Data is shown from a single flow rate curve for each fan speed (mean ± SD from at least 20 measurements of ΔCO2, recorded at 10 s intervals after each flow rate stabilized.

### Measurement of wind speed and boundary layer conductance inside the LI-6800F chamber

We next examined wind dynamics inside the LI-6800F chamber (**Fig. 2**). Gas exchange analysis revealed that fan speed increments from 0 up to 1000 rpm had a minor effect on wind speed (change from 0.005 to 0.029 m s^-1^), while a change in fan speed from 1000 to 10000 rpm increased wind speed linearly (at a slope of 0.0002 m s^-1^ per rpm) reaching 2 m s^-1^ at 10000 rpm. To further clarify the effect of g_b_ on transpiration, while eliminating the confounding effect of g_s_, we measured the transpiration rate from a wet filter paper under dark conditions. The response of boundary layer conductance (g_b filter paper_) was divided into 2 linear sections. First, at fan speeds from 300 rpm to 3100 rpm g_b filter paper_ increased from 0.16 to 1.42 mol m^-2^ s^-1^ (slope of 0.0005 mol m^-2^ s^-1^ per rpm), and second from 3100 to 10000 rpm g_b filter paper_ increased up to 2.4 mol m^-2^ s^-1^ (slope of 0.0001 mol m^-2^ s^-1^ per rpm). It’s important to note that some discrepancies exist between our estimation of the boundary layer conductance and the conductance calculated by the LI-6800F at fan speeds lower than 1900 rpm and higher than 4000 rpm. We therefore replaced the values of the boundary layer conductance in the LI-6800F data sheet used to calculate *g*_*s*_ with the data obtained by the filter paper method in our measurements. In addition to estimating g_b_ using the filter paper we calculated the width of the boundary layer in the 6800F 2 cm^2^ chamber according the to the relationship:

**Figure 2:**
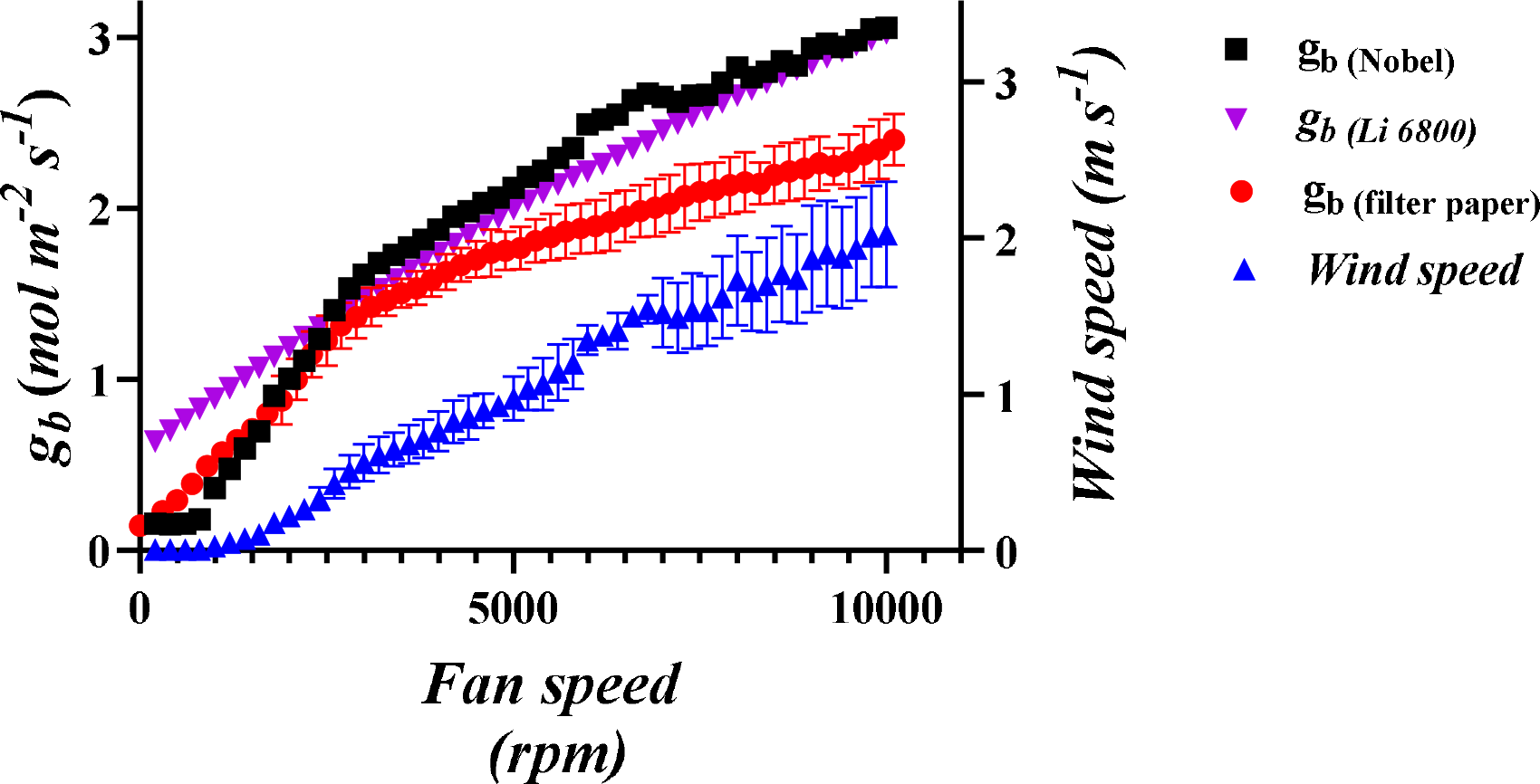
Effect of fan speed on wind speed and boundary layer conductance inside the LI-6800 chamber. Wind speed was measured using an omnidirectional hot wire sensor and boundary layer conductance was estimated by the wet filter paper method. The measurements were conducted under dark conditions, a filter paper temperature (T_exchange_) of 22°C, and a leaf vapor pressure deficit (VPD) ranging between 0.6-1 kPa. The black squares represent boundary layer conductance calculated following the methodology outlined by Nobel at each wind speed (2020). The purple triangles represent the boundary layer conductance calculated by the Licor software at each fan speed. The red circles represent the boundary layer conductance of the filter paper (mean ± SD from four separate determinations), estimated as the total leaf conductance calculated by the LI-6800 (gtw). The blue triangles represent the wind speed at the leaf plane inside the chamber at each fan speed ((mean ± SD from nine separate measurements).

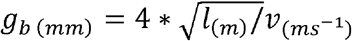 (Equation 7.10 Nobel 2009) where l_(m)_ is the diameter of the chamber, v _(_m s^-1^ is the ambient wind speed at each fan speed as measured by us, and g_b (mm)_ is the thickness of the boundary layer in mm. Next, we used the relationship g_b_ = D^w^ / g_b (mm)_ (Aphalo & Jarvis, 1993), where D^w^ is the diffusion coefficient of water in air to calculate the approximate *g*_*b*_ in mol m^2^ s^-1^ for all wind speeds measured inside the chamber from fan speed of 200 rpm to 10000 rpm. The calculated data agreed well with our estimation only at the lower range of fan speeds from 800 to 2700 rpm and with the original data calculated by the LI-6800 only in the higher part of the range above 2700 rpm (black line Fig. 1). At the very low range (<800 rpm) the calculation diverged from both our measurements and the LI-6800 model.

### Manipulating wind speeds to modulate transpiration rates and vapor pressure deficit

Changing the wind speed from 0.03 to 2 m s^-1^ resulted in *g*_*s*_ increasing from 0.33 to 0.51 mol m^-2^ s^-1^ within one minute in Arabidopsis (**Fig 3a**). Our results indicate that by manipulating fan speeds from 1000 to 10000 rpm we could induce changes in E (from 2.2 to 3.1 mmol m^-2^ s^-1^) with an inverse effect on VPD (0.9 to 0.68 kPa) due to the cooling effect of the increased transpiration. It important to note that the reverse effect (reduction in g_s_ in response to lower fan speed) was also measured (**Fig S2**). These results demonstrate that higher g_s_ is linked to elevated E resulting from enhanced g_b_.

**Figure 3:**
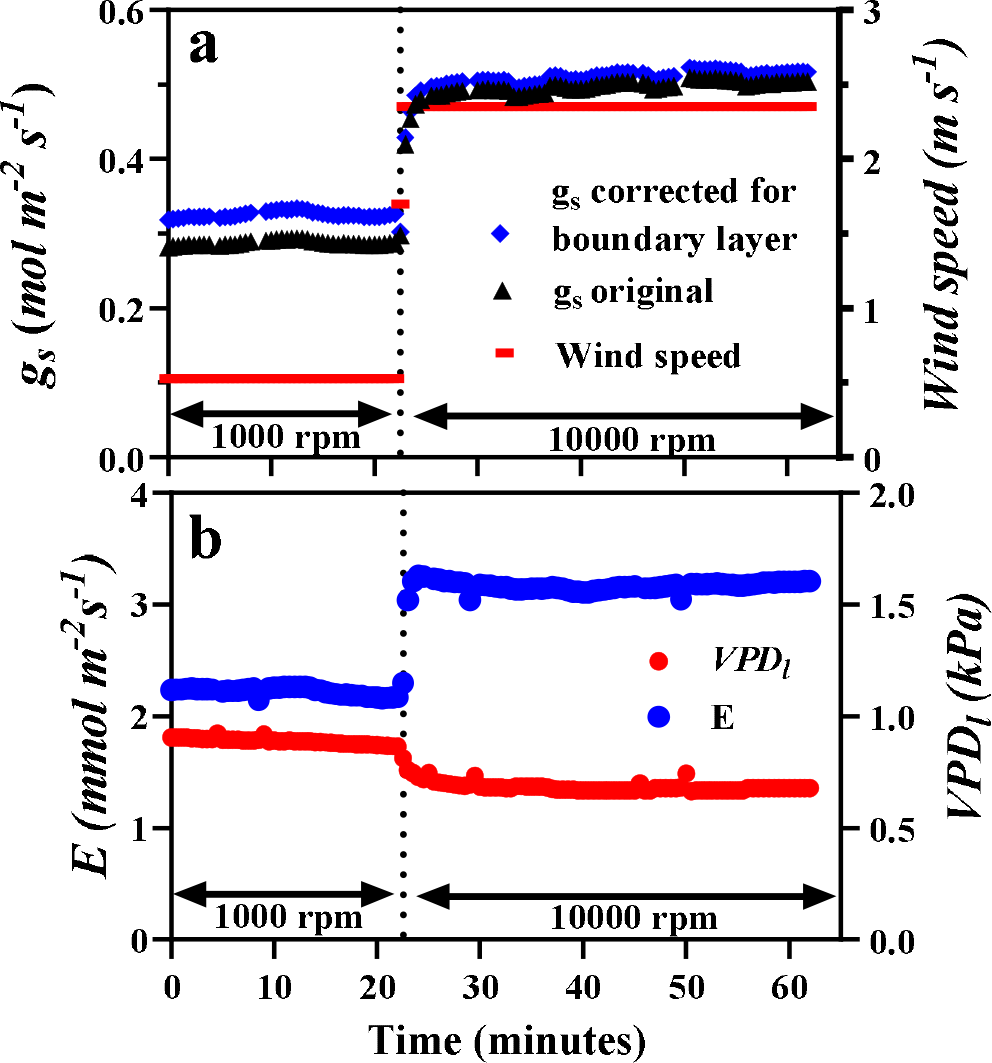
The effect of a rapid increase from low (1000 rpm) to high (10000 rpm) fan speed on Arabidopsis: (a) stomatal conductance (g_s_) according to the boundary layer estimated by the LI-6800 (black triangles) and after correcting the boundary layer conductance according to our measurements (blue circles), and wind speed (red squares), and (b) transpiration (E, blue circles) and leaf-to-air vapor pressure deficit (VPD_l_, red circles). Measurements were carried out at a CO_2_ concentration of 100 ppm to minimize the effect of internal CO_2_ concentration on stomatal aperture.

A gradual increase in wind speed over 5 min, ranging from 0.005 (fan speed of 200 rpm) to 1.5 m s^-1^ (fan speed of 7000 rpm) (**Fig. 4a**) corresponded to a progressive rise in g_s_ from 0.2 to 0.33 mol m^-2^ s^-1^. The gradual increment in wind speed resulted in an increase in E from 1.4 to 2.7 mmol m^-2^ s^-1^, accompanied by a decrease in VPD_l_ from 1.2 to 0.95 kPa (**Fig. 4b**). The fact that the increase in E actually changed *g*_*s*_ is demonstrated by the change in photosynthesis rate, which increased from 8.2 to 11.2 µmol m^-2^ s^-1^ over the 5-min interval of wind speed increment (**Fig. 4c**). It is worth noting that the transpiration efficiency decreases by 31%, suggesting a more pronounced impact on water vapor relative to assimilation in response to an increase in wind speed.

**Figure 4:**
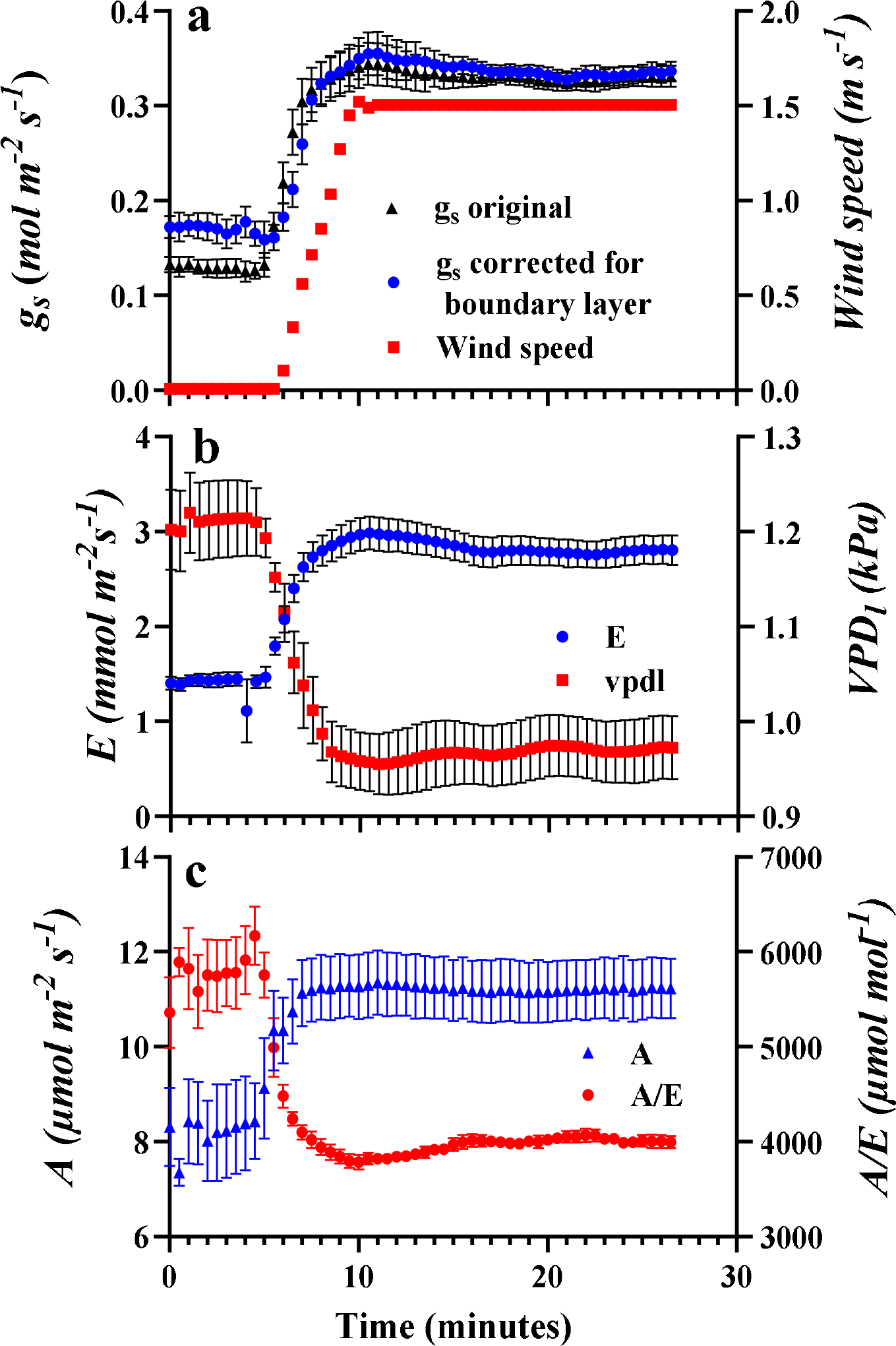
The effect of gradual changes in wind speed on Arabidopsis: (a) stomatal conductance (g_s_) according to the boundary layer estimated by the LI-6800 (black triangles) and after correcting the boundary layer values according to our measurements (blue circles), (b) transpiration (E, blue circles) and leaf-to-air vapor pressure deficit (VPD_l_, red squares), and (c) photosynthesis (A, blue triangles) and transpiration efficiency (A/E, red circles) at ambient CO_2_ (415 ppm). Wind speed was changed by adjusting the leaf fan speed from 200 rpm to 7000 rpm over a period of five minutes. Data shown as mean ± SE from four independent replications.

### Stomatal conductance response to light under different fan speeds

We examined stomatal opening kinetics in the transition from dark to light in *Vicia faba*. We compared three different fan speeds with stomatal opening observed in epidermal peels (**Fig. 5**). We found a significant relationship between fan speed and the rate of stomatal opening. As fan speed increased, the rate of stomatal opening also increased, suggesting that higher g_b_ is associated with faster stomatal opening kinetics. Stomatal opening observed in epidermal peels was similar to the opening seen during gas exchange measurements with fan speeds of 500 rpm, but 14 and 63% slower compared to stomatal opening during gas exchange measurements with fan speeds of 1000 and 5000 rpm, respectively.

**Figure 5:**
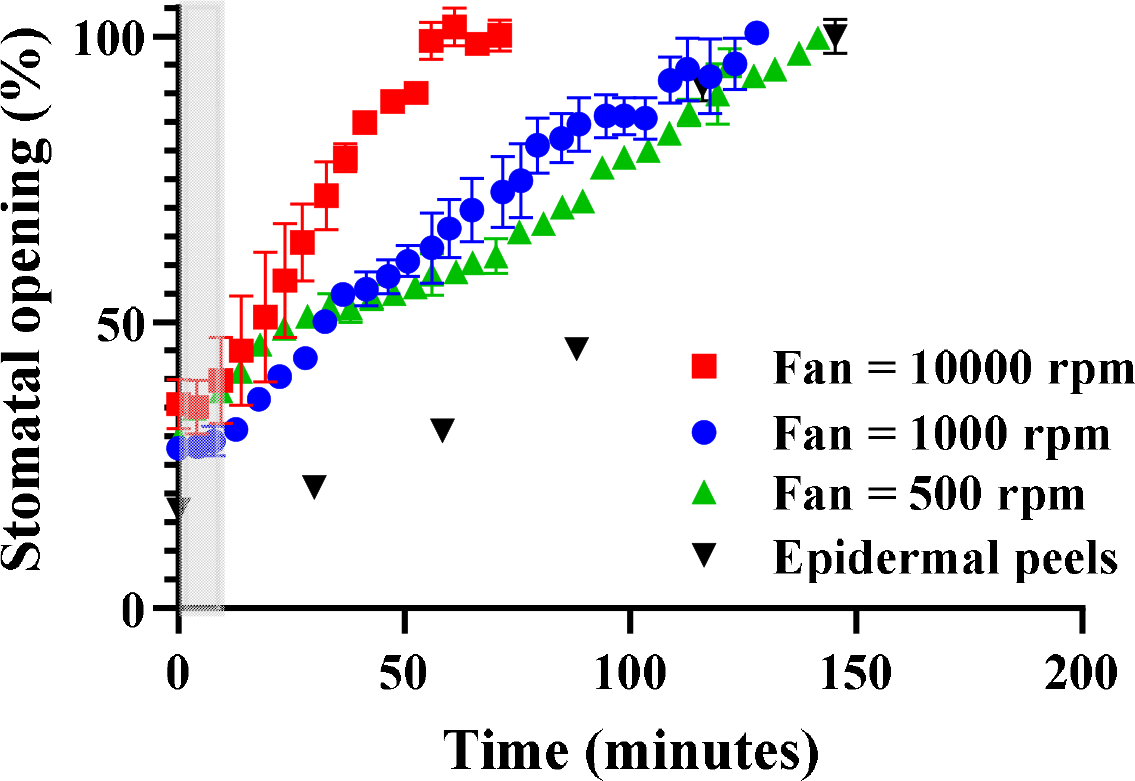
Changes in stomatal conductance throughout the transition from darkness to light (800 µmol m^-2^ s^-1^) for leaves of *Vicia faba* subjected to fan speeds of 500, 1000, and 10000 rpm. Additionally, the data illustrates variation in stomatal aperture observed in epidermal peels in a buffer solution derived from leaves collected from the same plants. The data are expressed as a percentage of stomatal opening [(g_s max_-g_s_)/ g_s max_] to facilitate comparison between gas exchange and epidermal peel measurements. Error bars represent the standard deviation from three independent measurements (n=3).

### Stomatal conductance response to increasing wind speed in various plant species

We investigated the response of g_s_ to increasing wind speed in several plant species growing under field conditions, including mango (*Mangifera indica*), papaya (*Carica papaya*), *Withania somnifera* and fig (*Ficus carica*) (**Fig. 6**). We found a consistent enhancement in g_s_ for all species as wind speed increased from 0.05 to 2.5 m s^-1^. The inset in figure 6 shows the slopes of the regression fits with their 95% CI. These findings highlight the impact of wind speed on stomatal behavior across the different plant species studied.

**Figure 6:**
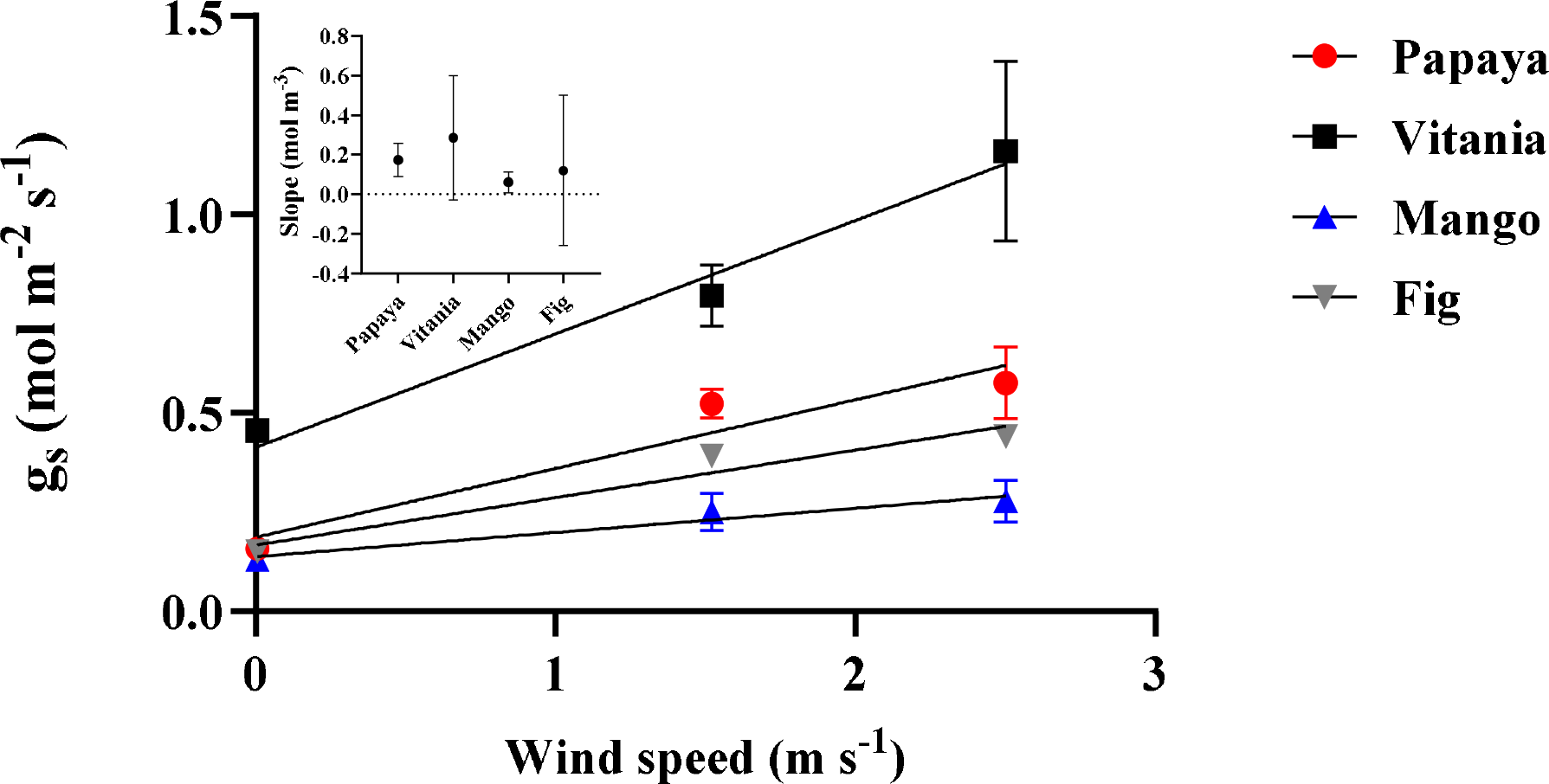
The effect of wind speed within the LI-6800 chamber on stomatal conductance in multiple plant species (corrected for boundary layer based on filter paper). Error bars represent the standard deviation from three independent measurements (n=3). Inset shows the linear slopes of the regression fit between g_s_ and wind speed ± 95% CI.

## Discussion

### Technical consideration for gas exchange measurements at low fan speeds

Our results indicate that the LI-6800F gas exchange system’s air mixing capability is not compromised even under fan speed as low as 200 rpm and is mainly dependent on the air flow rate rather than fan speed (**Fig. 1**). This flow-driven mixing should not come as a surprise if we consider that an airflow rate above 500 µmol s^-1^ in a chamber volume of 87 cm^3^, represents 12 complete air replacements every minute (1.1 L min^-1^). It thus seems that the recommendation to set the leaf fan to 10000 rpm (Using the LI-6800 v2.1 https://www.licor.com/env/support/LI-6800/manuals.html) is not critical for accurate measurements of gas exchange. The ability to measure gas exchange at low fan speed, combined with the versatile possibilities of fan speed control provided by new gas exchange systems, opens a range of possibilities for studies of leaf response to wind.

We found that the filter paper estimation of g_b_ did not always agree with g_b_ according to Nobel, (2020) and was lower especially at fan speed above 2700 rpm (**Fig. 2**). One reason that our g_b_ data was lower could be related to the pattern of the air flow across the leaf surface in the chamber. The model proposed by Nobel centers the boundary thickness calculation on the laminar flow of air where air movement is predominantly parallel to the leaf surface. In the gas exchange chamber air movement cannot be absolutely parallel to the leaf surface due to the geometry of the chamber air inlets. The leaf surfaces are sunk below the leaf gasket most likely preventing the conditions required for the Nobel equation calculation. Another deviation could result from the discrepancy between the mean length of the leaf in the direction of the wind (in the model *l*_(m)_) and the *l*_(m)_ which we used as the diameter of the 2 cm^2^ round chamber. Sub-saturation of the filter paper leading to lower transpiration and interpreted as lower total conductance (g_*t*_) could also be a reason for the discrepancy between our results and Nobel, (2020).

More importantly, our estimation of *g*_*b*_ diverged from the data supplied by the manufacturer (**Fig. 2**). We are not sure regarding the source of this error, but we would like to clarify that it probably makes little significance with respect to past measurements. The vast majority of published measurements were made at max fan speed (e.g., Barzilai et al., 2021; Sperling et al., 2014; Zait et al., 2019) meaning that g_b_ was high. Because g_s_ is significantly smaller than g_b_ and since resistances are summed 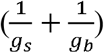, there is little impact of inaccuracy in g_b_ when resolving g_s_ under high fan speed. For example, when measuring a leaf with a g_s_ of 0.2 mol m^-2^ s^-1^, changing g_b_ from 2 mol m^-2^ s^-1^ to 3 mol m^-2^ s^-2^ would increase the overall conductance from 0.182 to 0.188 mol m^-2^ s^-1^. The estimation of g_b_ becomes far more important under low fan speed, when inaccuracy in the g_b_ model could result in a significant impact on g_s_ estimation (see difference between the blue and black lines in **Figs. 3** and **4**). It is thus critical to accurately estimate g_b_ when measuring at low fan speeds.

### The effect of wind on stomatal mechanics and opening kinetics

The effect of wind on stomatal conductance has important implications for determining stomatal kinetics, understanding stomatal mechanics, and quantifying stomatal regulation in response to environmental variables. It has long been known that the epidermal cells of angiosperms interact with the guard cells to determine stomatal aperture (Darwin, 1898; Iwanoff, 1928) and several studies demonstrated that equal loss of turgor in both guard and epidermal cells result in stomatal opening (Glinka, 1971; Franks *et al*., 1998; Franks & Farquhar, 2007). Our results suggest that this guard cell and epidermal cell mechanical interaction influences the dynamics of stomatal responses to both light and evaporative demand.

The most well-described manifestation of the epidermal interaction with the guard cells is the wrong-way opening of stomata in response to leaf excision and rapid dehydration (Franks & Farquhar, 2007; Zait et al., 2017). This counter-intuitive response, wherein higher conductance transiently occurs as leaf water status declines, can also occur at high VPD (Buckley et al., 2011). The opening is thought to occur passively due to increase in transpiration, which leads to lower turgor of both guard cells and epidermal cells and a corresponding increase in stomatal aperture due to the mechanical advantage of epidermal cells (ref). Our results, which showed a similar effect of increased g_b_ to that of increased VPD_l_, indicates that it is increased transpiration that drives the rapid passive stomatal response to leaf water status via epidermal turgor, rather than direct signaling induced by relative humidity or temperature (**Fig. 3**). This point is further reinforced by the fact the VPD_l_ decrease with the increase in wind speed is reversible without hysteresis (**Fig. 4, S2**). Our results align with those of Mott et al (1990) showed that increased transpiration due to exposing leaves to helox gas (2.3 times higher vapor diffusion relative to air) also resulted in significant stomatal opening. This line of evidence suggests that the wrong-way stomatal responses due to changes in transpiration are a mechanism by which angiosperms can passively regulate stomata. However, the passive responses of angiosperm stomata are in the opposite direction to the passive, “right-way” hydraulic regulation of stomatal responses to changes in leaf water status observed in lycophytes and ferns (Brodribb and McAdam 2011).

From a quantitative perspective our data shows that relatively mild increases in transpiration (from 2.2 to 3.2 mmol m^-2^ s^-1^) results in a significant stomatal opening (from 0.3 to 0.5 mmol m^-2^ s^-1^). Since such transpiration increase can be driven by common environmental changes (e.g. VPD increase from 1 to 2 kPa, a fairly common change for leaves that transition from shade to sunlight, is expected to double transpiration) This highlights that many of the current stomatal regulation models, which couples the extent and rate of stomatal movement only to ion transport, are incomplete (Jezek et al., 2019). Jezek et al. (2019) found that the stomatal opening kinetics predicted by OnGuard models (focus on solute transport) were three-to-five times slower than in vivo gas exchange observations. No parameter adjustments within physiological ranges brought the model kinetics significantly closer to experimental data, indicating a missing component in the model construction. The model prediction is in line with the stomatal opening that we have documented in epidermal peels, highlighting that transpiration, and its passive effect on epidermal turgor, could be the missing component. We suggest that integrating the transpiration effect on stomatal regulation, as influenced by wind (or VPD) (**Fig. 7**) should improve the model’s ability to predict the actual kinetics of stomatal movement.

**Figure 7:**
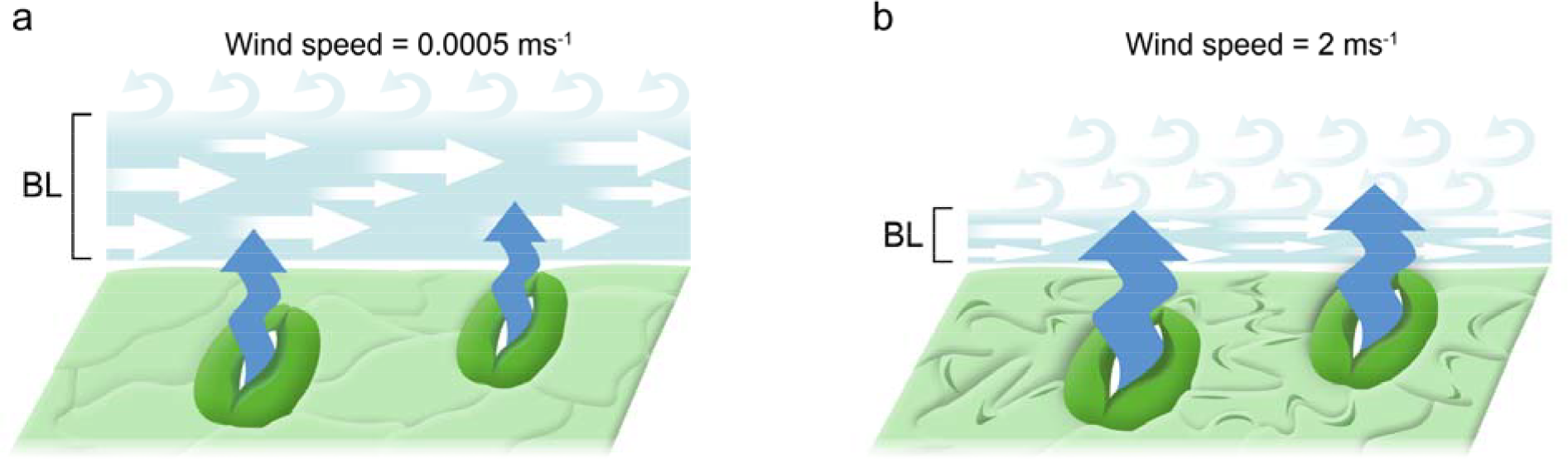
**(a)** Illustration of low wind speed scenario. The boundary layer (BL) around the leaf is relatively thick, impeding the diffusion of water vapor from the leaf surface to the external environment, thereby slowing transpiration (represented by thinner blue arrows coming out of the stomatal pores). **(b)** Condition of high wind speed. In this case, the increased air movement disrupts and thus thins the boundary layer around the leaf. A thinner boundary layer facilitates the diffusion of water vapor from the leaf surface to the external environment, accelerating transpiration (indicated by thicker blue arrows). The increase in transpiration creates a greater water deficit in the leaf tissues, leading to a decrease in turgor pressure within the epidermal cells, especially those surrounding the stomata. This loss of turgor pressure passively causes the stomata to open, facilitating further transpiration and allowing greater uptake of CO_2_.

It should be noted that most past studies that exposed plants to higher wind speed found that they exhibit lower g_s_ (e.g. Renard & Demessemacker, 1983). This is probably the result of unfavorable hydration conditions due to the higher transpiration, similar to the reduced g_s_ of plants under high VPD (reference). This is not necessarily the case when a single leaf is placed inside the gas exchange cuvette. The increased transpiration from a single leaf, has little impact on whole plant water use and a negligible effect on xylem water potential. Accordingly, the transpiration of a well hydrated leaf can be increased to some extent before it will lead to either epidermal turgor loss, or loss of mesophyll cell turgor and the triggering of ABA biosynthesis that might drive active stomatal closure (McAdam & Brodribb, 2016).

### Considerations for accurate g_s_ measurements

The fact that wind speed has such a dramatic effect on g_s_ measurements, raises questions about our ability to accurately estimate the native g_s._ The effect is expected to be most pronounced when leaves are taken from low-wind environments and placed in a gas exchange cuvette with high fan speed that rapidly increase their transpiration and open their stomata, meaning that the recorded value is higher than the native g_s_. Most past studies have used high fan speed because this is the official recommendation of gas exchange manufacturers, but it is a far more complicated challenge to understand the wind speed that individual leaves experience in their natural environment. Multiple studies report that the wind speed inside dense canopies of an agricultural crop or forests are significantly lower than those measured by the meteorological stations (typically located out of the field or forest; Shaw, 1977; Renaud et al., 2011). This suggests that measuring leaves in conditions above their ambient wind, and consequently inducing stomatal opening, is not rare. This notion is also supported by the fact that upscaling leaf gas exchange measurements into canopy scale typically result in overestimation of the whole plant transpiration (Flore, 2003; Hochberg et al., 2023).

Adjusting for potential artefacts requires careful consideration. Firstly, one should take into account the ambient wind speed and direction and, in combination with the leaf dimensions, determine the native g_b_ as suggested by Nobel (2020). Subsequently, adjusting the cuvette fan speed helps in reproducing a similar boundary layer. However, it’s noteworthy that in many instances, users might be looking to assess gas exchange under standardized conditions rather than the ambient (manifested in the common procedure to use a controlled temperature,CO_2_ concentration, RH and fan speed).

To conclude, wind speed can have a large effect on stomatal conductance and stomatal kinetics. Incorporating this effect into the current stomatal dogma and hydraulic models should improve our ability to predict plant response to the environment. Accounting for the wind effect is critical when measuring leaves using gas exchange system.

## Acknowledgment

This study supported by the Israel Science Foundation (ISF) grant 2076/23 to Y.Z.

## References

Aphalo, P. J., & Jarvis, P. G. (1993). The boundary layer and the apparent responses of stomatal conductance to wind speed and to the mole fractions of CO2 and water vapour in the air. Plant, Cell & Environment, 16(7), 771–783. 10.1111/j.1365-3040.1993.tb00499.x

Assmann, S. M., & Jegla, T. (2016). Guard cell sensory systems: recent insights on stomatal responses to light, abscisic acid, and CO2. Current Opinion in Plant Biology, 33, 157–167. 10.1016/j.pbi.2016.07.003

Barzilai, O., Avraham, M., Sorek, Y., Zemach, H., Dag, A., & Hochberg, U. (2021). Productivity versus drought adaptation in olive leaves: Comparison of water relations in a modern versus a traditional cultivar. Physiologia Plantarum, 173(4), 2298–2306. 10.1111/ppl.13580

Buckley, T. N. (2005). The control of stomata by water balance. New Phytologist, 168(2), 275–292. 10.1111/j.1469-8137.2005.01543.x

Buckley, T. N., Sack, L., & Gilbert, M. E. (2011). The role of bundle sheath extensions and life form in stomatal responses to leaf water status. Plant Physiology, 156(2), 962–973. 10.1104/pp.111.175638

Cowan, I. R. (1978). Stomatal Behaviour and Environment. Advances in Botanical Research, 4(C), 117–228. 10.1016/S0065-2296(08)60370-5

Darwin, F. (1898). Observations on stomata. Transactions of the Royal Society of London, 58(1496), 212–213. 10.1038/058212a0

DeMichele, D. W., & Sharpe, P. J. H. (1973). An analysis of the mechanics of guard cell motion. Journal of Theoretical Biology, 41(1), 77–96. 10.1016/0022-5193(73)90190-2

Flore, G. F. & J. A. (2003). Comparison Between Different Methods for Measuring Transpiration in Potted Apple Trees. Biologia Plantarum, 46(1), 41–47.

Foster, J. R., & Smith, W. K. (1986). Influence of stomatal distribution on transpiration in lowLwind environments. Plant, Cell & Environment, 9(9), 751–759. 10.1111/j.1365-3040.1986.tb02108.x

Franks, P. J. (2013). Passive and active stomatal control: Either or both? New Phytologist, 198(2), 325–327. 10.1111/nph.12228

Franks, P. J., Cowan, I. R., & Farquhar, G. D. (1998). A study of stomatal mechanics using the cell pressure probe. Plant, Cell and Environment, 21(1), 94–100. 10.1046/j.1365-3040.1998.00248.x

Franks, P. J., & Farquhar, G. D. (2007). The mechanical diversity of stomata and its significance in gas-exchange control. Plant Physiology, 143(1), 78–87. 10.1104/pp.106.089367

Frensch, J., & Schulze, E. D. (1988). The effect of humidity and light on cellular water relations and diffusion conductance of leaves of Tradescantia virginiana L. Planta, 173(4), 554–562. 10.1007/BF00958969

Glinka, Z. (1971). The Effect of Epidermal Cell Water Potential on Stomatal Response to Illumination of Leaf Discs of Vicia faba. Physiologia Plantarum, 24(3), 476–479. 10.1111/j.1399-3054.1971.tb03521.x

Hochberg, U., Perry, A., Rachmilevitch, S., Ben-Gal, A., & Sperling, O. (2023). Instantaneous and lasting effects of drought on grapevine water use. Agricultural and Forest Meteorology, 338(May), 109521. 10.1016/j.agrformet.2023.109521

Iwanoff, L. (1928). Zur Methodik der Transpirationsbestimmung am Standort. Ber Dtsch Bot Ges, 46:, 306–310. 10.1007/BF00419279

Jezek, M., Hills, A., Blatt, M. R., & Lew, V. L. (2019). A constraint–relaxation–recovery mechanism for stomatal dynamics. Plant Cell and Environment, 42(8), 2399–2410. 10.1111/pce.13568

Kearns, E. V., & Assmann, S. M. (1993). The guard cell-environment connection. Plant Physiology, 102(3), 711–715. 10.1104/pp.102.3.711

Kim, T.-H., Böhmer, M., Hu, H., Nishimura, N., & Schroeder, J. I. (2010). Guard Cell Signal Transduction Network: Advances in Understanding Abscisic Acid, CO2, and Ca2+ Signaling. Annual Review of Plant Biology, 61(1), 561–591. 10.1146/annurev-arplant-042809-112226

Kollist, H., Nuhkat, M., & Roelfsema, M. R. G. (2014). Closing gaps: Linking elements that control stomatal movement. New Phytologist. 10.1111/nph.12832

Mcadam, S. A. M., & Brodribb, T. J. (2016). Linking Turgor with ABA BiosynthesislJ: Implications for Stomatal Responses to Vapor Pressure De fi cit across Land Plants 1 [ OPEN ]. 171(July), 2008–2016. 10.1104/pp.16.00380

McAdam, S. a M., & Brodribb, T. J. (2014). Separating active and passive influences on stomatal control of transpiration. Plant Physiology, 164(4), 1578–1586. 10.1104/pp.113.231944

McNab, B. K. (2006). The relationship among flow rate, chamber volume and calculated rate of metabolism in vertebrate respirometry. Comparative Biochemistry and Physiology - A Molecular and Integrative Physiology, 145(3), 287–294. 10.1016/j.cbpa.2006.02.024

Meidner, H., & Heath, O. V. S. (1963). Rapid changes in transpiration in plants. In Nature (Vol. 200, Issue 4903, pp. 283–284). 10.1038/200283a0

Mott, K. a., & Franks, P. J. (2001). The role of epidermal turgor in stomatal interactions following a local perturbation in humidity. Plant, Cell and Environment, 24(6), 657–662. 10.1046/j.0016-8025.2001.00705.x

Nobel, P. S. (2020). Physicochemical and environmental plant physiology. In Physicochemical and Environmental Plant Physiology (5th ed.). Academic Press. 10.1016/C2018-0-04662-9

Parkinson, K. J. (1985). A simple method for determining the boundary layer resistance in leaf cuvettes. Plant, Cell & Environment, 8(3), 223–226. 10.1111/1365-3040.ep11604618

Raschke, K. (1970). Leaf Hydraulic System: Rapid Epidermal and Stomatal Responses to Changes in Water Supply. Science, 167, 189–191.

Renard, C., & Demessemacker, W. (1983). Effects of wind velocity on stomatal conductance and consequences of leaf rolling on water uptake in tall fescue. Biologia Plantarum, 25(6), 408–411. 10.1007/BF02903136

Renaud, V., Innes, J. L., Dobbertin, M., & Rebetez, M. (2011). Comparison between open-site and below-canopy climatic conditions in Switzerland for different types of forests over 10 years (1998-2007). Theoretical and Applied Climatology, 105(1), 119–127. 10.1007/s00704-010-0361-0

Shaw, R. H. (1977). Secondary Wind Speed Maxima Inside Plant Canopies. Journal of Applied Meteorology and Climatology, 16(May), 514–521.

Shimazaki, K., Doi, M., Assmann, S. M., & Kinoshita, T. (2007). Light regulation of stomatal movement. Annual Review of Plant Biology, 58, 219–247. 10.1146/annurev.arplant.57.032905.105434

Sperling, O., Lazarovitch, N., Schwartz, A., & Shapira, O. (2014). Effects of high salinity irrigation on growth, gas-exchange, and photoprotection in date palms (Phoenix dactylifera L., cv. Medjool). Environmental and Experimental Botany, 99, 100–109. 10.1016/j.envexpbot.2013.10.014

Zait, Y., Shapira, O., & Schwartz, A. (2017). The effect of blue light on stomatal oscillations and leaf turgor pressure in banana leaves. Plant, Cell & Environment, 40(January), 1143–1152. 10.1111/pce.12907

Zait, Y., Shtein, I., & Schwartz, A. (2019). Long-term acclimation to drought, salinity and temperature in the thermophilic tree Ziziphus spina-christi: Revealing different tradeoffs between mesophyll and stomatal conductance. Tree Physiology, 39(5), 701–716. 10.1093/treephys/tpy133

Zhu, M., Jeon, B. W., Geng, S., Yu, Y., Balmant, K., & Chen, S. (2016). Preparation of Epidermal Peels and Guard Cell Protoplasts for Cellular, Electrophysiological, and -Omics Assays of Guard Cell Function. Methods in Molecular Biology, 1363(November), 89–121. 10.1007/978-1-4939-3115-6_9.

