## Supplemental for "Wind speed affects the rate and kinetics of stomatal conductance"

**Supplemental materials:**

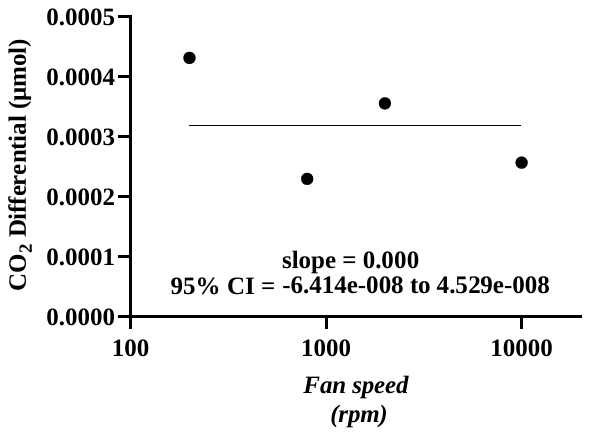

**Figure S1:** Slopes of the regression between CO_2_ differential and flow rate at 4 different fan speeds (200, 800 2000 and 10000 rpm) were compared using linear regression. The slopes were found to be similar with the regression slope between them and the fan speed is determined by a symmetric 95% CI around zero.

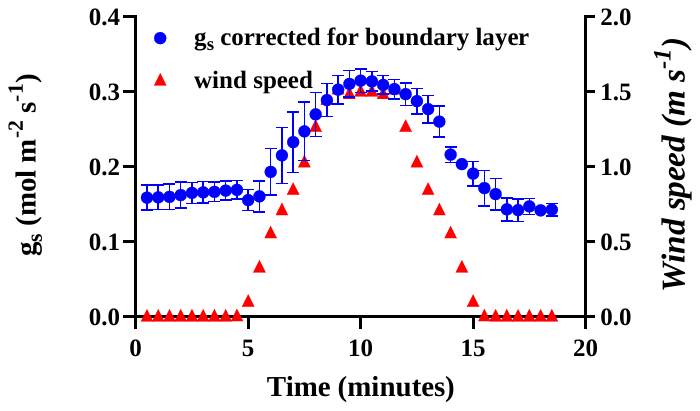

**Figure S2:** Response of stomatal conductance to a gradual increase and decrease of wind speed (mean ± SE, n=4).

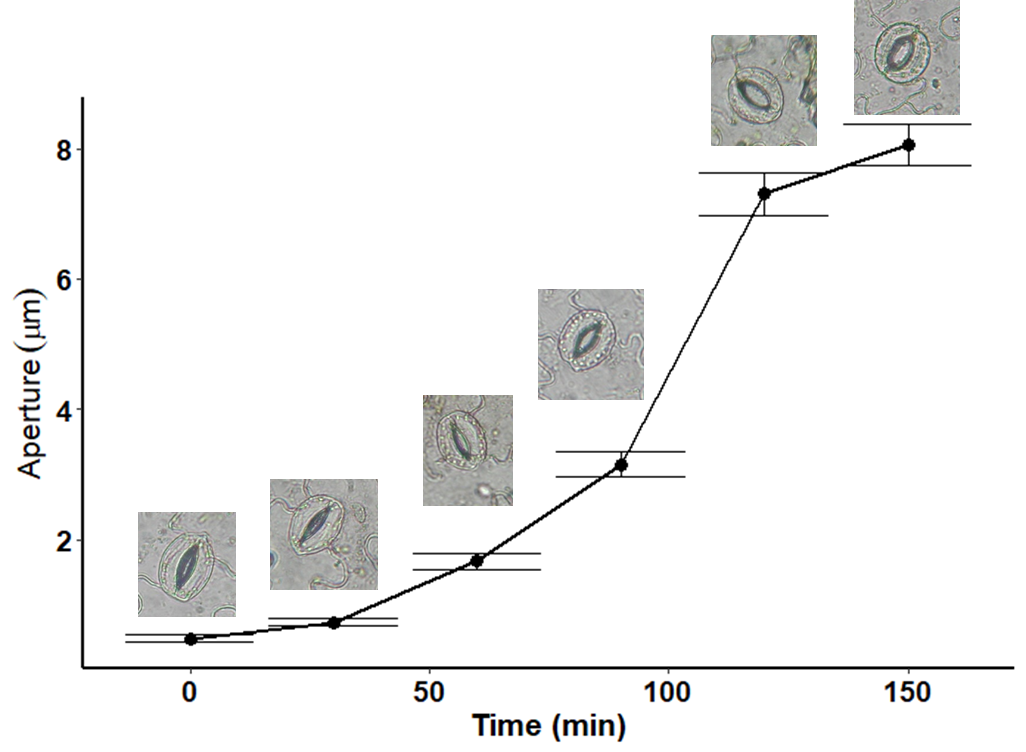

**Figure S3:** Time course of stomatal aperture in epidermal peels of *Vicia faba* during the transition from darkness to light (400 µmol m^-2^ s^-1^). Stomatal aperture was measured at regular intervals following the introduction of light. The error bars in the graph represent the standard error of three plants, with measurements taken from a total of 30 stomata per peel.
